# Detecting Material State Changes in the Nucleolus by Label-free Digital Holographic Microscopy

**DOI:** 10.1101/2023.03.17.533098

**Authors:** Christiane Zorbas, Aynur Soenmez, Jean Léger, Christophe De Vleeschouwer, Denis L.J. Lafontaine

## Abstract

Ribosome biogenesis is initiated in the nucleolus, a multiphase biomolecular condensate formed by liquid-liquid phase separation. The nucleolus is a powerful disease biomarker and stress biosensor whose morphology reflects its function. Here we have used digital holographic microscopy (DHM), a label-free quantitative phase contrast microscopy technique, to detect nucleoli in adherent and suspension cells. Subtle nucleolar alterations induced by drug treatment or by depletion of ribosomal proteins were efficiently detected by DHM. We trained convolutional neural networks to detect and quantify nucleoli automatically on DHM images of cultured human cells (HeLa). Holograms containing cell optical thickness information allowed us to define a novel nucleolar index which we used to distinguish nucleoli whose material state had been optogenetically modulated. We conclude that DHM is a powerful tool for quantitatively characterizing nucleoli, including material state, without any staining.

## INTRODUCTION

Ribosomes are essential nanomachines responsible for protein production in cells. Ribosome biogenesis is initiated in the nucleolus, a dynamic biomolecular condensate formed by liquid-liquid phase separation (LLPS) (Lafontaine *et al*, 2020). The nucleolus is the most prominent membraneless organelle in the cell nucleus, whose morphology reflects its role in ribosome biogenesis and other functions important for cell homeostasis (Boisvert *et al*, 2007; Pederson, 1998). The nucleolus is rich in RNA and proteins that associate transiently through multivalent weak interactions to perform functions (Feric *et al*, 2016; Mitrea *et al*, 2016). The dynamic nature of the nucleolus can be explained in part by its liquid properties. These can be optogenetically tuned, with an impact on function (Zhu *et al*, 2019).

The nucleolus is a potent disease biomarker and stress biosensor, as its size, shape, and number per cell nucleus are markedly altered in cancer, viral infections, neurodegeneration, and ageing (Boulon *et al*, 2010; Derenzini *et al*, 2009; Salvetti & Greco, 2014; Tiku & Antebi, 2018). Despite its remarkable properties, the nucleolus remains largely underused in clinical work for lack of easy-to-implement, robust quantitative tools.

Nucleolar structure has been abundantly studied at the light and electron microscopy levels for many years (Hernandez-Verdun *et al*, 2010). In human cells, the nucleolus is organized in three major layers, nested like Russian dolls, and encased in a sheath of perinucleolar chromatin. The three main internal layers are the fibrillar center (FC), the dense fibrillar component (DFC), and the granular component (GC). A single GC contains multiple modules, each comprising an FC core surrounded by a DFC.

A wide range of microscopy techniques have been used in combination with dedicated staining methods to detect the nucleolus quantitatively in cultured cells and tissue biopsies. In principle, this requires either particular labeling chemistry, for example silver nitrate-based AgNOR staining of the argyrophilic proteins which abound in the nucleolus (Bartholome *et al*, 2019; Ploton *et al*, 1986; Thelen *et al*, 2021), the use of specific antibodies for immunodetection, or expression of fluorescently tagged proteins for direct detection (Nicolas *et al*, 2016; Stamatopoulou *et al*, 2018; Stenstrom *et al*, 2020). More recently, super-resolution techniques have been applied to fixed and live samples, revealing the existence of additional nucleolar subphases (Ide *et al*, 2020; Yao *et al*, 2019). While these techniques are extremely powerful in describing the most intricate details of this fascinating biomolecular condensate, their high sophistication remains a clear obstacle to their routine use.

Digital holographic microscopy (DHM), invented by Dennis Gabor in 1948 (Gabor, 1948), is a non-invasive label-free quantitative interferometric technique that can be applied with minimal manipulation to any transparent specimen, such as fixed or live cells. Unlike most microscopy techniques, which record absorption and transmission of light from an object, DHM records shifts in the light wavefront with respect to a reference wavefront, producing a digital hologram from which a phase image is extracted digitally by numerical reconstruction (**Fig 1**). A key feature distinguishing DHM from optical-contrast-enhancing imaging techniques is that in DHM, the intensity value of a pixel has a direct physical meaning: it is proportional to the *optical thickness* (also called *optical path length*) of the cell, which is the physical height of the cell (or cell thickness) multiplied by its refractive index at that point (Picart, 2015; Picart & Li, 2012). Because of well-described optical artifacts, such quantitative information cannot be extracted from the images obtained by conventional brightfield microscopy or other contrast-enhancing imaging techniques such as Zernicke’s’ phase-contrast (PhC) microscopy or Smith and Nomarski’s differential interference contrast (DIC) microscopy (see (Marquet *et al*, 2005) for details). DHM captures can be displayed as intuitive pseudo 3-D images resembling topographic maps, where the height is determined by the brightness of each pixel (**Fig 1** and **Fig 2A**). Thus far, DHM has been exploited in material science, cell biology, and cancer studies. It has notably been used to monitor cell structure and dynamics in various biological and biomedical contexts, such as cell growth monitoring, cell dry mass estimation (Barer, 1952), drug-induced cytoskeleton dynamics (Kemper *et al*, 2006), neuronal growth, and metastasis progression (for a review see (Marquet *et al*, 2014). DHM is also particularly well suited for imaging liquid samples and performing in-flow analyses (Singh *et al*, 2017).

**Figure 1.**
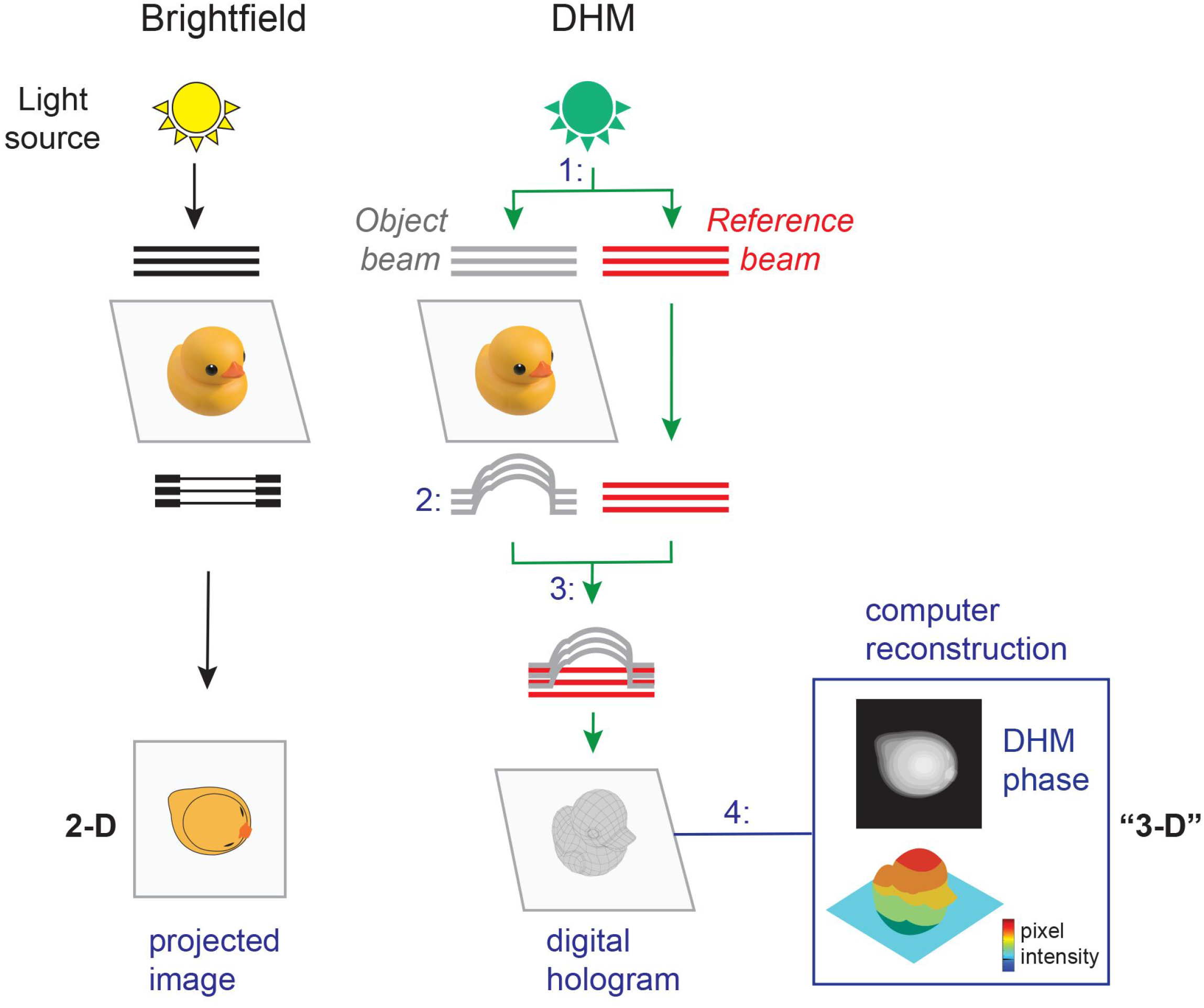
Principle of digital holographic microscopy (DHM). **A** In brightfield microscopy, a light source illuminates an object (here illustrated as a duck), dense areas of the specimen absorb light, and the transmitted light is projected and recorded as an image. **B** In DHM, in order to generate an ‘interference pattern’ or hologram, 1: an illumination source is split into an ‘object beam’ (object wave light) and a ‘reference beam’ (reference wave light), 2: the object beam passes through the sample and is subjected to a phase shift, creating the ‘object wave front’, 3: the ‘object wave front’ and the ‘reference wave front’ are combined to interfere and to create a hologram which is captured with a CCD camera, 4: a numerical reconstruction algorithm is used to produce a phase image from the digitally captured hologram. The phase image, which represents the optical thickness of the object at each point, can be displayed as a pseudo 3-D map using pixel intensity.

**Figure 2.**
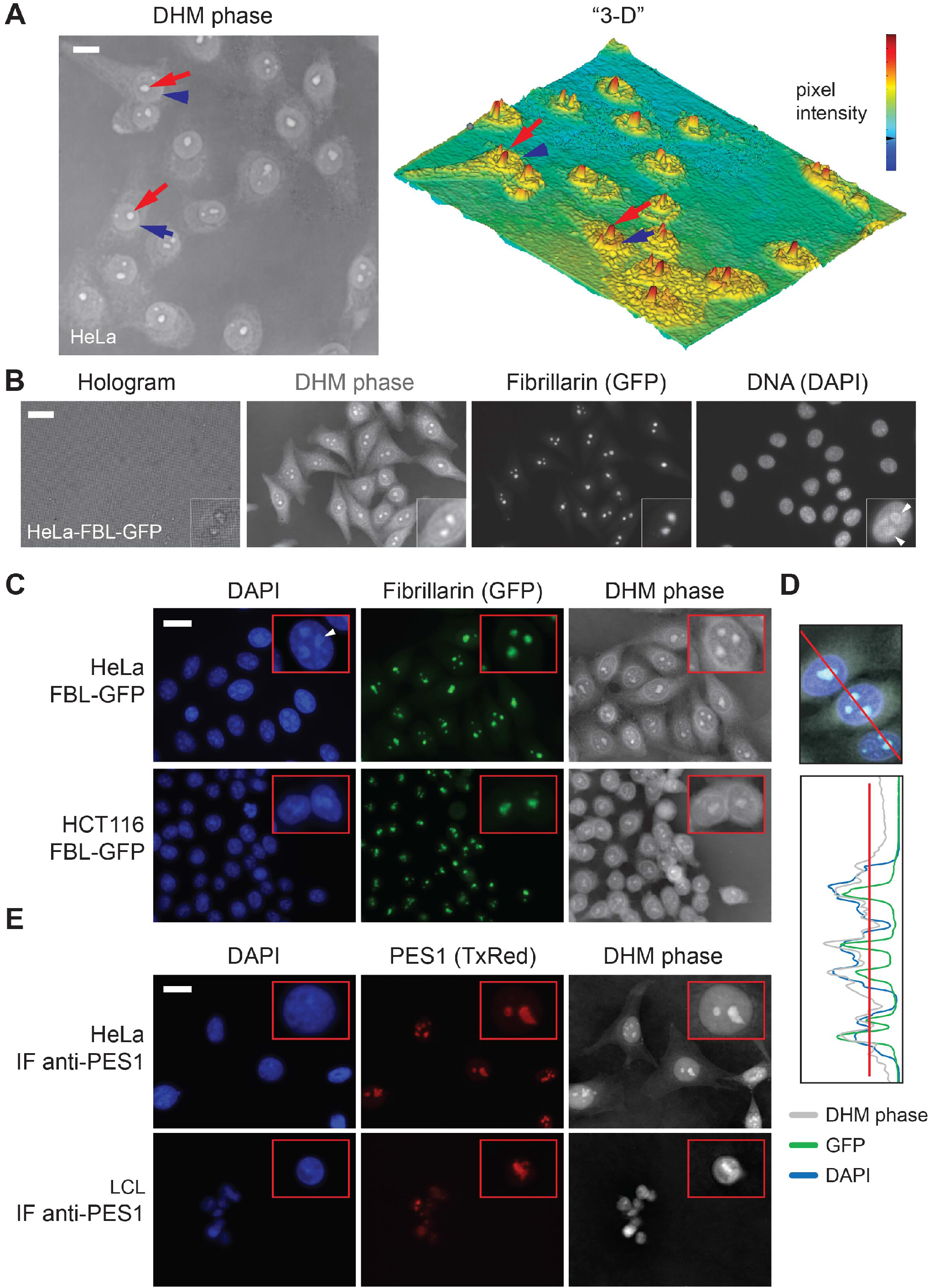
The nucleolus can be detected by DHM. **A** (Left) HeLa cells visualized by DHM. In each cell, the contour of the nucleus is readily detectable (blue arrowhead); within the nucleus one or several intense structures are detected (red arrow); these correspond to nucleoli. Scale bar, 20 μm. (Right) Pseudo 3-D map display of the image shown on the left, based on pixel intensity representing the optical thickness. **B** HeLa cells stably expressing the green-fluorescently-tagged nucleolar protein fibrillarin (HeLa-FBL-GFP) observed by correlative DHM-fluorescence microscopy. Cells were stained with DAPI, which labels the DNA-rich nucleoplasm. The hologram, DHM phase, green fluorescence (GFP), and DAPI signals are shown. Insets, magnification of an individual cell nucleus, illustrating the perinucleolar chromatin ring (arrowheads in the DAPI panel). Scale bar, 20 μm. **C** HeLa-FBL-GFP and HCT116-FBL-GFP cells observed by correlative DHM-fluorescence microscopy. The perinucleolar chromatin ring is also visible in the DAPI channel (white arrowhead). **D** Quantification traces of the DHM phase (in gray), green fluorescence (in green), and DAPI (in blue) signals. **E** Immunodetection of the nucleolar protein PES1 in HeLa and LCL cells visualized by correlative DHM-fluorescence microscopy. PES1 was imaged in red (Texas red, TxRed). In panels C and E, the DAPI, green fluorescence, and DHM phase signals are shown. Insets, magnification of an individual cell nucleus. Scale bar, 20 μm.

Here we have used DHM imaging of human cells combined with deep learning to detect and characterize the nucleolus quantitatively without any staining. The numerical parameters extracted include the mean number of nucleoli per cell nucleus, the mean nucleolar area, the mean nucleolar-to-nuclear area ratio, and the *nucleolar optical thickness*, a novel index which detects material states in the nucleolus.

## RESULTS & DISCUSSION

### Detection of the nucleolus by digital holographic microscopy

Unlike other phase-contrast-generating techniques and brightfield, digital holographic microscopy is quantitative, because in a DHM phase image the intensity of each pixel reflects the optical thickness of the cell, i.e. its physical height multiplied by its refractive index (Picart, 2015; Picart & Li, 2012). With this in mind, we expected optically dense cell structures, such as the nucleolus, to appear on DHM phase images as bright spots, and, less dense objects, such as vacuoles or the cytoplasm, as less intense areas. As shown below, this is indeed the case.

To test if DHM is suitable for detecting nucleoli, various cell lines were observed, starting with a common model: cervix carcinoma cells (HeLa). Cells were grown in a channel slide and observed directly under the microscope after brief fixation. It was also possible to observe live cells (see below). The nucleus contour was clearly identifiable in all cells (**Fig 2A**, blue arrowheads). Within the nucleus, prominent structures reminiscent of nucleoli were detected (red arrows). From the quantitative information embedded in the DHM phase image, a “3-D” map was generated from pixel intensities, revealing the optical thicknesses of individual cell substructures (**Fig 2A**, right). In such maps, the structures reminiscent of nucleoli appeared as sharp red peaks.

In our initial work, we used a standard digital holographic microscope with a beam path exactly as described in **Figure 1** (e.g. the phase image and 3-D display in **Fig 2A**). Soon we realized that if we wanted our method to be used in other laboratories worldwide, it would be greatly advantageous to use a “plug-in” device that would convert any inverted microscope to DHM. In the remainder of our work, we used a purposely built DHM adaptor (QMOD), also referred to as ‘off-axis differential interferometer’, directly connected to a classical CCD camera and to an inverted microscope (see **Fig EV1**). In this easy-to-implement set up (see Materials and Methods), the beam path was slightly more complex than that described in **Figure 1**, but the principle of quantitative interferometry was exactly the same.

A major advantage of the QMOD is that DHM can be readily combined with fluorescence imaging to perform correlative DHM-fluorescence microcopy. This is what we did to ascertain that the prominent structures detected in the nucleus were indeed nucleoli (**Fig 2B-E**). Initially, we used HeLa cells stably expressing the nucleolar protein fibrillarin fused to a green fluorescent tag (HeLa-FBL-GFP). The nucleolus consists of three main layers nested like Russian dolls (Thiry & Lafontaine, 2005), and fibrillarin marks the middle layer or dense fibrillar component. Cells were stained with DAPI, which labels the DNA-rich nucleoplasm. Comparing the fluorescence signal (fibrillarin, GFP) with the DHM phase made it obvious that the prominent nuclear substructures detected by DHM were nucleoli. The nucleolus is lined by a layer of condensed chromatin, the so-called perinucleolar chromatin forming a distinctive DNA “ring” around the nucleolus. This ring was visible in the DAPI images (see arrowheads in **Fig 2B**, DAPI inset). The presence of DNA rings circling the prominent nuclear foci observed in the DHM phase images further confirmed their identity as nucleoli. Quantification of the fluorescence and DHM signals demonstrated an excellent overlap between phase intensity and GFP peaks, formally confirming colocalization (**Fig 2D**).

To extend our observations to other cells, we used a colon carcinoma cell line (HCT116), also stably expressing a FBL-GFP construct (**Fig 2C**). In these cells, the DHM phase images again revealed prominent signals in the nucleus, again confirmed to be nucleoli on the basis of colocalization with fibrillarin and counter staining with DAPI (**Fig 2C**).

In addition to using fluorescently tagged nucleolar proteins, we detected endogenous proteins by indirect immunofluorescence with specific antibodies. In this experiment, we used both adherent cells (HeLa) and suspension cells (lymphoblastoid cells, LCL) and chose to detect the nucleolar protein PES1 (pescadillo ribosomal biogenesis factor 1, **Fig 2E**). PES1 labels the cortical layer of the nucleolus or granular component. In both cases, we observed excellent colocalization between the DHM phase and the fluorescence signal (**Fig 2E**). This again confirmed that the prominent nuclear foci detected by DHM were nucleoli.

Finally, four additional cell lines were observed: another type of cervix carcinoma cells (SiHa), bone cancer cells (U2OS), breast cancer cells (MCF7) and lung cancer cells (A549) (**Fig EV2**). All cells displayed prominent nucleolar signals, similar to those observed in HeLa and HCT116 cells.

In conclusion, DHM allows robust stain-free detection of the nucleolus in adherent and suspension cells of various tissue origins.

### Nucleolar structure alterations are detectable by digital holographic microscopy

Morphological alterations of the nucleolus are associated with diverse pathological conditions ranging from cancer, viral infection, and neurodegeneration to various types of cell stress and even ageing (Boulon *et al*., 2010; Derenzini *et al*., 2009; Tiku & Antebi, 2018).

We wondered if such nucleolar alterations could be detected by DHM. To test this possibility, we induced nucleolar alterations by treating cells with an array of drugs known to disrupt the nucleolus (Burger *et al*, 2010) or by depleting them of factors important for nucleolar structure maintenance (Nicolas *et al*., 2016; Stamatopoulou *et al*., 2018). We treated HeLa cells stably expressing an FBL-GFP construct with either DRB (5,6-dichloro-1-β-D-ribofuranosyl-1H-benzimidazole), roscovitine, actinomycin D, or CX-5461, or depleted them of ribosomal protein uL5 (formerly RPL11) or uL18 (RPL5) (**Fig 3**). Using correlative DHM-fluorescence microscopy, we detected fibrillarin and found each treatment to affect nucleolar structure deeply, as previously described in the literature (**Fig 3**, GFP panels). Comparing the fluorescence and DHM phase signals revealed that each alteration was perfectly detectable by DHM (**Fig 3**). Considering the absence of staining, the definition of nucleolar granules was truly exceptional in DHM phase images (see e.g. DRB images).

**Figure 3.**
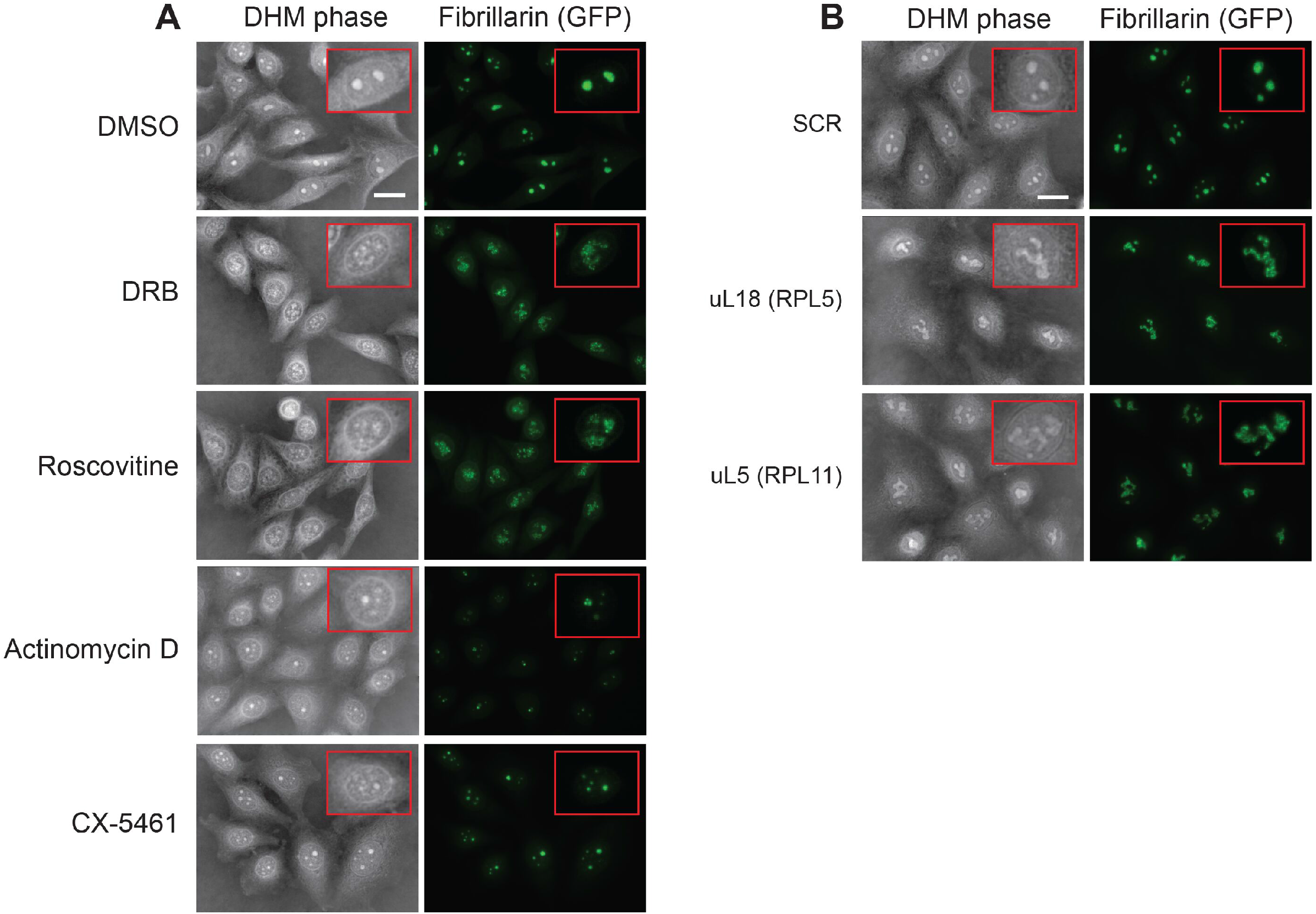
Nucleolar alterations can be detected by DHM. Diverse nucleolar alterations were induced in HeLa-FBL-GFP cells and observed by correlative DHM-fluorescence microscopy. **A** Drug-induced nucleolar alterations. Cells were treated with the vehicle control (DMSO 0.1%), DRB (32 μg/mL), roscovitine (50 μM), actinomycin D (0.2 μg/mL), or CX-5461 (5 μM) for 2 hours. Scale bar, 20 μm. **B** Nucleolar alterations induced by ribosomal protein knockdown. HeLa-FBL-GFP cells were treated for 3 days with an siRNA (10 nM) specific to the mRNA encoding ribosomal protein uL18 or uL5 or with a non-targeting scramble control (SCR). Scale bar, 20 μm.

In conclusion, DHM can readily detect fine alterations of nucleolar structure induced by pharmacological treatment or by depletion of ribosomal proteins important for nucleolar structure maintenance.

The nucleolus is visible only during the interphase, being known to disassemble at the onset and reassemble at the end of mitosis (Lafontaine *et al*., 2020). We wanted to know if nucleolar disassembly and reassembly could be monitored by DHM. We thus performed live-cell imaging using DHM and concluded that it can (**Movie 1**).

As discussed above, nucleolar alterations induced by drug treatments can be visualized efficiently by DHM on fixed cells (**Fig 3**). To see if such drug-induced alterations could also be monitored dynamically in live cells, we seeded HeLa-FBL-GFP cells into channel slides, added roscovitine, and imaged live cells every 30 seconds for 3 hours. The drug-induced nucleolar alterations could indeed be followed in live cells by DHM, since the changes observed in the GFP channel were also obvious in the phase signal (**Movie 2**). Although not the subject of our study, another interesting observation emerged from the movies: we saw numerous cytoplasmic granules, presumably mRNA granules such as processing bodies, exploring rapidly the cytoplasmic space. This indicates that the technology is suitable for studying other biomolecular condensates.

### Quantitative analysis of the nucleolus detected by DHM, using deep learning

Having shown that the nucleolus can be detected by DHM, we next sought to use the technique to extract types of quantitative information that would normally require specific staining. Typically, numerical parameters such as the mean number of nucleoli per cell nucleus, the mean nucleolar area, and the mean nucleolar-to-nuclear area ratio have proved useful in clinical biology.

A database of seventy-five fields of view was generated, each comprising about fifteen HeLa cells expressing fluorescently tagged fibrillarin. For each field of view, three images were captured: GFP, DAPI, and the DHM phase. This database was used to establish a fluorescent thresholding method for detecting nuclei and nucleoli and to train U-net convolutional neural networks. Importantly, all segmentation procedures were carefully benchmarked manually.

First, by means of fluorescence only, the nucleus and nucleoli of each cell were segmented, respectively, by thresholding the DAPI and GFP signals (**Figs 4A, EV3A, EV5A**). For detection of the nucleus, a simple thresholding method was sufficient, while for counting the nucleoli and establishing the nucleolar area, a more sophisticated thresholding method was required. This involved detecting local maxima, followed by ‘region growing’ (see Materials and Methods). During this thresholding, most cells undergoing mitosis (during which the nucleolus is disassembled, see (Lafontaine *et al*., 2020)) and multi-nucleated or incomplete cells (image edges) were filtered out. Importantly, only structures whose GFP signal was contained within a DAPI signal were considered to be nucleoli. The nucleolar parameters extracted are presented in **Figure 4B**. Altogether, 2882 nucleoli from 1206 cells were analyzed. This led to the conclusion that the mean number of nucleoli per HeLa cell was 2.39, the mean nucleolar area was 23.15 μm^2^, and the mean nucleolar-to-nuclear area was 0.14.

**Figure 4.**
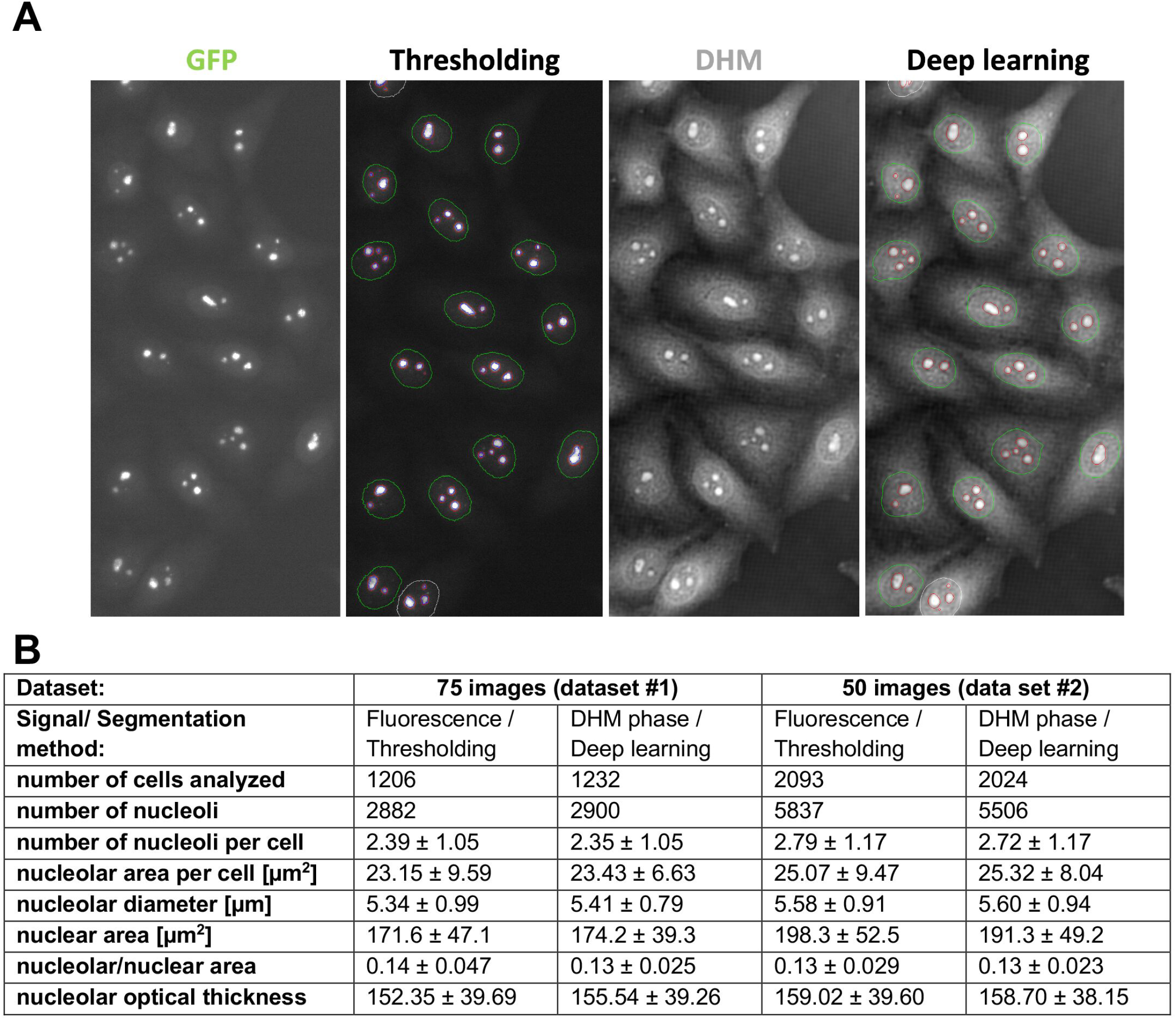
Nucleolar parameter analysis. **A** Illustration of image segmentation of HeLa cells stably expressing fibrillarin-GFP. The GFP channel was segmented by thresholding and the DHM phase by deep learning. On the left figure: red, automatic segmentation of nucleoli on GFP images by ‘region growing’, for computing the nucleolar area; blue: automatic segmentation of nucleoli by ‘region growing’, for counting; green: automatic segmentation of the nuclei by thresholding. On the right figure: red, automatic segmentation of nucleoli (for counting, area computation, and nucleolar optical thickness computation); green, automatic segmentation of the nuclei. **B** Data analysis. The parameters were extracted automatically by thresholding (fluorescence) or deep learning (DHM phase). The nucleolar optical thickness was calculated as the mean DHM intensity of a fixed area. To benchmark the robustness of our approach, data captured more than two years apart by two different scientists were compared (75-image set and 50-image set).

Interestingly, the mean nucleolar area was reasonably well conserved in nuclei containing up to four nucleoli (**Fig EV4A**), after which it gradually increased. This was as expected if small nucleoli coalesce into larger ones in liquid-liquid like fashion, in agreement with the LLPS model of nucleolar assembly. If one views the nucleolus as a sphere, the projected areas of multiple small spheres cover a larger area than the projection of fewer large spheres. Assuming that a constant volume V is divided into N identical spheres, the radius of each sphere becomes proportional to (V/N)^1/3^. Hence, the surface projected by N spheres is proportional to N times (V/N)^2/3^, which is proportional to N^1/3^ (**Fig EV4A**). Also note that automatic counting of nucleoli on fluorescence images was carefully benchmarked with manual annotations and assessed with confusion matrices as well as sensitivity, specificity, and precision scores (**Figs EV4B-C**). The fraction of cells whose nucleoli were accurately counted (i.e., the precision) was superior to 83% for cells displaying one to four nucleoli. The sensitivity and specificity were >78% and >94%, respectively, when 1-4 nucleoli were counted. The most frequent errors were counting one too many nucleoli or missing one. The count was less accurate when five or more nucleoli were counted, but this situation was hardly ever encountered in cells (**Fig EV4B**).

The next step was to identify the nucleus and nucleoli in cells directly on the DHM phase images, without using fluorescence signals. To achieve this, two 2D U-net convolutional neural networks were trained (Falk *et al*, 2019), one using as input the segmented DAPI signal, the other using the segmented GFP signal (**Figs 4, EV3B-C, EV5B**). The database of images was divided into three groups of twenty-five fields of view each. Two groups were combined for training and the third was used for testing. The operation was reiterated twice until each group had been used once for testing. All seventy-five fields of view were then used for automatic extraction of numerical features. As an illustration, a representative GFP image was segmented by thresholding and the corresponding DHM phase by deep learning, with nearly identical results (**Figs 4A, EV5**). The data show that the numerical parameters extracted directly from the DHM phase images by deep learning were highly consistent with those obtained by thresholding the GFP images (**Fig 4B**). The histograms representing the number of nucleoli per cell nucleus computed from the fluorescence images and those computed from the DHM phase images are nearly identical (**Fig EV4B**). The precision, sensitivity, and specificity computed from the confusion matrix were respectively >64%, >64%, and >80%, when one, two, or three nucleoli were counted (**Fig EV4C**).

As discussed in the Introduction, pixel intensity has a biological meaning in DHM phase images. On this basis, we introduced the ‘nucleolar optical thickness’ as a novel parameter corresponding to the mean nucleolar DHM intensity (**Fig 4B**). For HeLa cells, the nucleolar optical thickness was ~150. Precisely, it was 155 when extracted directly from the phase images by deep-learning-based segmentation, and 152 when binary masks generated by fluorescence thresholding were applied to DHM phase images (**Fig 4B**).

Once established this new method for extracting quantitative nucleolar parameters, it was important to test its robustness. An independent dataset based on 50 new images was produced. The two datasets were acquired by independent scientists more than two years apart. The comparative analysis revealed high consistency of the extracted parameters (**Fig 4B**). The differences observed may reflect marginal metabolic fluctuations associated with cell passage number, medium batch, etc.

### Changes in the material state of the nucleolus can be detected by DHM

At this stage, we wanted to explore the potential use of the newly defined nucleolar optical thickness. With recent increased interest in cell biology for soft matter research and biophysics, it has become clear that the material state of a cell can be modulated locally, notably by use of optogenetics, with an impact on function. We were particularly interested in learning if DHM might be suitable for assessing different material states of the nucleolus.

To test this idea, we designed an experiment where we converted the nucleolus from a liquid to a gel by use of optogenetics (Zhu *et al*., 2019). Briefly, we engineered a Cry2olig tag-containing nucleolar construct and expressed it directly from the genome of an HEK293 cell. The Cry2olig tag is known to self-polymerize upon exposure to blue light (488 nm), this leading to protein aggregation and a change in material state (**Fig 5A-B**)(Taslimi *et al*, 2014; Zhu *et al*., 2019). To ascertain nucleolar targeting and efficient mixing of the Cry2olig fusion protein with the nucleolar phase, a nucleolar localization signal (NoLS) was inserted into the construct in addition to an intrinsically disordered domain (IDR). Additionally, an mCherry tag was used to monitor subcellular distribution by fluorescence microscopy and protein mobility by fluorescence recovery after photobleaching (FRAP).

**Figure 5.**
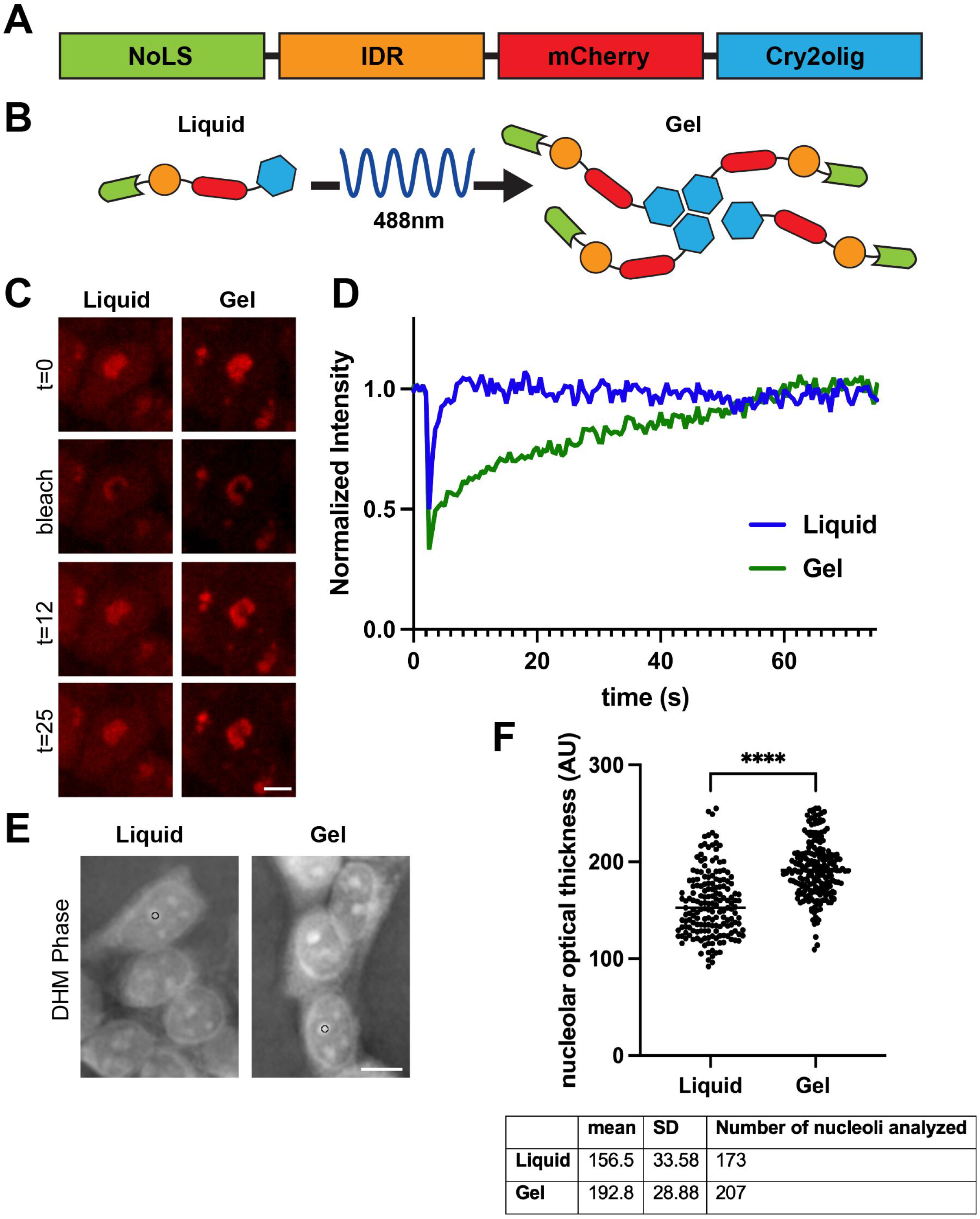
Material state alterations can be detected by DHM. **A** Structure of the Opto-tag construct used. NoLS, nucleolar localization signal; IDR, intrinsically disordered region; mCherry, fluorescent tag for microscopy; Cry2olig; self-polymerization tag activated upon blue light exposure (see Materials and Methods for details). **B** Rationale of material state change: upon exposure to blue light, the opto-tag construct self-polymerizes, turning the nucleolus from a liquid to a gel. **C, D** Fluorescence recovery after photobleaching (FRAP) in control cells (liquid state) and cells exposed to blue light (gel). Time scale (t), secs. FRAP analysis was conducted on the same cells before and after blue-light-induced protein aggregation. Panel C presents an example of a photobleached cell and panel D, the matching FRAP curves. **E** DHM phase images of cells prior to and after exposure to blue light. Black circle, a constant disk area defined for quantification (12 pixels). **F** Quantification of optical thickness in DHM phase images. Three hundred and eighty nucleoli were quantified. Nucleolar gelation led to a 23% increase in the nucleolar optical thickness index. ****, Mann-Whitney test, p<0.0001.

Upon exposure to blue light, the mobility of the Cry2olig protein construct decreased, confirming gelation of the nucleolus (**Fig 5C-D**). We used DHM to compare the nucleolar optical thicknesses of 380 gelated and non-gelated nucleoli. In cells exposed to blue light, we observed a significant 23% increase in nucleolar optical thickness. In conclusion, DHM can efficiently distinguish nucleolar material states (**Fig 5E-F**).

In summary, considering that the nucleolus is a powerful yet underused biomarker and biosensor, there is a need to develop robust, easy-to-implement quantitative detection techniques that typically do not involve staining. This is exactly what we have achieved in this work, by applying digital holographic microscopy. We show that in multiple adherent and suspension human cells, DHM detects prominent signals in the nucleus, corresponding to nucleoli (**Figs 2, EV2**). The nucleolar nature of the structures detected in cell nuclei was demonstrated by colocalization with nucleolar antigens by correlative DHM-fluorescence (**Fig 2C-E**). The nucleolar antigens used for colocalization were either fluorescently tagged proteins (fibrillarin) or endogenous proteins detected with specific antibodies (PES1). Two of the three main layers of the nucleolus were represented in the colocalization studies, as fibrillarin and PES1 respectively label its dense fibrillar and granular components. The nucleolar nature of the observed structures was further confirmed by the presence of a perinucleolar chromatin ring visible by DAPI counterstaining (**Fig 2B**). Very fine alterations of nucleolar structure, induced by pharmacological treatments or by depletion of factors important for ribosome biogenesis (such as specific ribosomal proteins) were also perfectly detectable by DHM (**Fig 3**).

To extract numerical features directly from DHM phase images, we have developed a deep learning strategy and have trained convolutional neural networks for automatic recognition of the nucleus and nucleolus. From analysis of our images, we conclude that in HeLa cells, the mean number of nucleoli per cell is ~2.5, the mean nucleolar surface is ~24 μm^2^, and the mean ratio of nucleolar-to-nuclear area is ~0.13 (**Fig 4**). Having extracted very similar numbers from our GFP images by thresholding and from our DHM images by deep learning, we conclude that our analysis pipeline is robust. The extracted features largely correspond to published values (Caragine *et al*, 2019; Farley *et al*, 2015; Puck *et al*, 1956), although in published data the sampling was never as deep as the 3,000 to nearly 6,000 nucleoli analyzed here all at once. This type of numerical parameters has high potential value in basic research on ribosome biogenesis and in mechanistic studies of processes as essential as tumorigenesis, viral infection, senescence, neurodegeneration, and ageing, among others where nucleolar morphology has been shown to vary greatly.

A unique aspect of DHM is that it provides information about optical thickness, which we used to define a novel nucleolar index. Reasoning that the novel index might be sensitive to the material state of the nucleolus, we have shown this to be the case, as nucleolar gelation was detected as a >23% increase in nucleolar optical thickness (**Fig 5**).

In conclusion, we have established digital holographic microscopy as a powerful method for characterizing the nucleolus numerically without any staining. We have introduced a novel parameter, the nucleolar optical thickness, and proved it to be useful in studying nucleolar material state changes. We believe DHM will be widely applicable to the study of numerous other biomolecular condensates of natural or artificial origin.

## MATERIALS AND METHODS

### Cell lines and cell culture

#### Cell lines used

All cell lines used in this work were purchased from ATCC and diagnosed by short tandem repeat (STR) profiling: HeLa (CCL-2), HCT116 (CCL-247), SiHa (HTB-35), MCF7 (HTB-22), U2OS (HTB-96), and A549 (CCL-185). HeLa-FBL-GFP cells have been described previously (Nicolas *et al*., 2016; Stamatopoulou *et al*., 2018). HCT116-FBL-GFP cells were generated by CRISPR-Cas9 editing, according to (Nakade *et al*, 2014). The LCL cell line is a kind gift from Dr Alyson W. MacInnes (Amsterdam UMC, The Netherlands).

#### Cell culture conditions

Cells were grown at 37°C under 5% CO_2_. HeLa, SiHa, and A549 cells were grown in DMEM (Lonza, BE12-604F) supplemented with 10% fetal bovine serum (Sigma, F7524), 1% penicillin-streptomycin mix (Lonza, DE17-602E). HCT116 and U2OS cells were grown in McCoy’s medium (Lonza, BE12-688F) supplemented with 10% fetal bovine serum, 1% penicillin-streptomycin mix. MCF7 cells were grown in EMEM (ATCC, 30-2003) supplemented with 10% fetal bovine serum, human recombinant insulin at 0.01 mg/mL (Sigma, I9278), and 1% penicillin-streptomycin mix. LCL cells were grown in RPMI (Lonza, BE12-167F) supplemented with 15% fetal bovine serum and 2 mM L-glutamine (Lonza, BE17-605E).

#### Cell preparation for DHM imaging

Unless otherwise stated, cells were grown in an Ibidi μ-slide I (Ibidi, 80106), fixed in methanol for 5 min at room temperature (RT), washed three times in 1x PBS (Lonza, 17-516F), incubated in DAPI (Sigma, D9542, 250 ng/ml, prepared in 1x PBS), and washed three times in 1x PBS for 5 min prior to imaging.

#### Correlative DHM-fluorescence detection

Cells were fixed in 2% formaldehyde (Sigma-Aldrich, F8775) for 15 min at RT, washed three times in 1x PBS for 5 min, permeabilized by incubation in 1x PBS /0.3% Triton X-100 (Sigma-Aldrich, X100)/5% BSA (Roche, 10735086001) for 1 hour at RT, incubated overnight at 4°C with an anti-PES1 antibody raised in rat (8E9, (Holzel *et al*, 2007)), and diluted 1:1000 in 1x PBS/0.3%Triton X-100/1% BSA, washed three times in 1x PBS for 5 min, incubated for 1 hour at RT with a goat antirat Alexa Fluor 594 antibody (A11007, Thermofisher Scientific) diluted 1:1000 in 1x PBS/0.3% Triton X-100/1% BSA, washed three times in 1x PBS for 5 min, incubated for 15 min at RT with DAPI (250 ng/ml, prepared in PBS 1x), and washed three times in 1x PBS prior to imaging.

#### Drug treatment

HeLa cells were grown overnight in Ibidi μslides and treated with the vehicle control (DMSO 0.1%, Sigma-Aldrich, D2650), roscovitine (50 μM, Sigma, R7772), DRB (32 μg/mL, Sigma, D1916), actinomycin D (0.2 μg/mL, Sigma A1410), or CX-5461 (5 μM), Sigma-Aldrich, 5092650001) for 2 hours prior to DHM imaging.

#### siRNA depletion

Cells were transfected with siRNAs as described in (Tafforeau *et al*, 2013). The siRNAs used (scramble, #4390843, RPL5, #s12153, and RPL11, #s12170) were purchased from Thermofisher Scientific.

### DHM Image capture

#### Hardware

Cells were observed by correlative DHM/fluorescence microscopy on a Zeiss Axio Observer Z1 driven by MetaXpress, equipped with LED illumination (CoolLed pE-2), and fitted with a QMOD off-axis differential interferometer (Ovizio s.a.) and a Retiga R3 camera (see Fig S1). Holograms were captured using transmitted light (HAL lamp fitted with a single-band bandpass optical filter (Semrock, FF01-550/49-25)) converted with a dedicated software routine. The DHM phase was produced with OsOne. Holograms were imaged with a 20x (0.5 NA) EC Plan Neofluar (Zeiss, 420350 – 9900) or a 40x (0.75 NA) EC Plan Neofluoar (Zeiss, 420360 – 9900) objective and converted to DHM phase with OsOne.

#### Pixel-to-size (μm) conversion

Image pixel size was determined as the camera pixel size multiplied by the binning divided by (objective magnification x lens magnification x C mount). In our set-up these values were: pixel size of the Retiga R3 camera, 4.54 x 4.54 μm; binning, 1x; objective magnification, 20x; lens magnification, 1.5x; C-mount, 1x. For capture at 20x magnification (Figure 2B and all the images used in the quantitative analysis), the image pixel size was (4.54 μm x 1)/(20 x 1.5 x 1)= 0,151 μm/pixel. Note that the pixel-to-size conversion was validated empirically with a calibration slide (Pyser-SGI, 02A00404).

### Image processing pipeline

#### Description of the database

The database was composed of two datasets of 75 and 50 triplets of images, respectively. A triplet consisted of aligned GFP, DAPI, and DHM images. The resolution of the GFP and DAPI images was 1460 x 1920 pixels and the resolution of the DHM images was 364 x 480. The DHM images were interpolated with a spline of order 3 so as to reach the same resolution as the GFP and DAPI images. All images were then cropped to 1408 x 1920, as the deep learning model used required every input image dimension to be a multiple of 2^5^. The nucleoli and nuclei were manually contoured on 25 triplets and 10 triplets on the first and second datasets, respectively. The two datasets were captured more than two years apart by two different scientists.

### Segmentation of the fluorescence images

#### Segmentation of the nucleoli on GFP images

The processing pipeline for segmenting the fluorescence images is depicted in Figure EV3A. The dynamic range of each GFP image was set at [0,1] by dividing the intensity of each pixel by its maximum intensity across the dataset. In order to lower the intensity variations across different images of the dataset, the GFP images were further standardized image-wise using the mean and standard deviation of the pixel intensities belonging to the nuclei in the considered GFP image (using the nuclei segmentation computed on the DAPI images). Then, nucleoli were identified as pixels whose intensity was above a certain threshold. A multistep approach was necessary because a single threshold did not allow both counting the number of nucleoli per cell nucleus and reliably measuring their area, as a too-low threshold led to fusion of neighbouring nucleoli while a too-high threshold led to underestimating the nucleolar area and loss of low-intensity nucleoli.

Therefore, our approach used a sequence of thresholds to first identify all local maxima in the GFP image, after which a region-growing algorithm was used (i) to merge local maxima connected by sufficiently intense pixels and (ii) to connect illuminated pixels to their closest local maximum.

The local maximum detection algorithm considers an increasing sequence of thresholds [*t*_0_, *t*_1_,…,*t_i_*, *t_N_*] ranging from *t*_0_ = 0.083 to *t_N_* = 0.3 with an increment of 0.025. The algorithm iterates from the lowest to the highest threshold. The connected regions segmented with *t*_*i*+1_ are always included in the regions obtained with *t_i_*. The algorithm allows a region to split into several subregions when the next threshold is considered. However, it does not allow a region to disappear. After each thresholding operation, two morphological operations are applied: an opening (using a disk with radius 2 as a structuring element) followed by a closing (using a disk with radius 3 as a structuring element). These operations force very close pixels to belong to the same region. When the highest threshold is reached, the seed of each final subregion is defined as its maximum. Hence, there are as many seeds as subregions at the last iteration of the algorithm.

Both region-growing algorithms expand from the computed seeds by adding adjacent pixels until a stopping pixel intensity criterion is met. The region-growing algorithms proposed for nucleolus counting and nucleolar area computation stop when the intensity of the added pixels reaches respectively *max*(*α* * *I_seed,j_, τ*) or *τ*, where *I_seed,j_* is the pixel intensity of seed *j* and *τ* is a constant threshold. Some cells in mitosis (which do not display well-formed nucleoli) are also excluded during this processing step. The GFP signal of those cells has a low intensity and a large area, in contrast to that of cells in interphase, which have either a bright GFP signal or a small area. Cells in mitosis are detected when a region-growing algorithm with a *T_mitosis_* stopping threshold provides a region with an area larger than 1000 pixels and a maximum intensity within the region smaller than 0.25. In those cases, the seed is ignored and no region is added. Both region-growing algorithms iterate on all the seeds. In some cases, small holes appear in the grown regions, which are filled. The parameters were tuned to *α* = 0.85, *τ* = 0.15 and *τ_mitosis_* = 0.11.

The outputs of automatic nucleolar segmentation are two black-and-white images (one for counting and the other for area computation) where white pixels belong to the nucleoli and black pixels to the background (Fig EV3A, see insets on the right).

#### Segmentation of nuclei on the DAPI images

The dynamic range of each DAPI image was set at [0,1] by dividing the intensity of each pixel by its maximum intensity. The threshold for segmenting DAPI images was set independently on every DAPI image with an automatic approach relying on the fact that the pixel intensity distribution is bimodal in DAPI images. The background and nucleus pixel intensities cluster, respectively, in the first and second modes of the distribution. Every DAPI image was filtered with a Gaussian kernel of standard deviation 3 and the histogram of pixel intensities was computed. Then the threshold was set at the abscissa value where the histogram reached a local minimum between its two modes. This heuristic showed better results than the Otsu algorithm (Otsu, 1979) on the considered dataset. A closing operation (using a disk with radius 3 as a structuring element) was applied in order to fill any small holes occurring within the segmented regions and to obtain a smoother nucleus contour. As isolated pixels could appear because of thresholding, all regions smaller than 2,000 pixels were removed in order to deal with those false detections. Then a dilation operation with a disk of radius 2 was applied to the binary mask obtained after the thresholding operation. The output of the automatic nucleus segmentation is a black-and-white image where white pixels belong to the nuclei and black pixels to the background (i.e., a binary mask, Fig EV3A, see inset). Finally, the binary masks for nucleoli and nuclei were multiplied element-wise in order to remove potential falsely detected nucleoli outside the nuclei in the automatic nucleolus segmentation outputs (see ⊗ symbol in Figure EV3A).

### Segmentation of the DHM images

#### Deep learning architecture

Nuclei and nucleoli were segmented directly on DHM images by means of deep learning. We used 2D U-net, a state-of-the-art deep learning architecture for biomedical image segmentation (Falk *et al*., 2019; Ronneberger *et al*, 2015). This architecture is a parameterized mathematical function (or model) that maps an input image to a desired output representation with the same size as the input image. In this study, the output representation is an image where the intensity of every pixel is defined between 0 and 1 by a sigmoid function, so that pixels that are close to one, i.e. above 0.5, are classified as a part of a nucleolus, whereas other pixels are classified as background pixels. The same approach is followed for nucleus segmentation. The inner parameters (called weights) of the deep learning model are adapted automatically through a training procedure in order to perform the desired task. This is performed by providing pixel-wise nucleolus or nucleus binary labels to the network in addition to the input images. The training phase proceeds iteratively, by randomly selecting small subsets of samples in the training dataset and adapting the neural network inner weights so that the output predictions fit the desired labels. Once a model is trained, it is used to segment new images, unseen during the training phase. During this testing phase, only input images are provided to the network, which then outputs a binary image with segmentation of the nucleolus or nucleus.

2D U-net belongs to a category of deep learning architectures called convolutional neural networks. These networks are particularly suited for image analysis, as a convolution operation acts as a filter that extracts image patterns. Simple filters can extract edges or corners, but when multiple filters are stacked, complex patterns can be recognized in images. Convolutional neural networks are essentially stacking filters organized in so-called convolutional layers. The 2D U-net model also uses maxpooling layers, in order to merge semantically similar features into one, and rectified linear units (ReLu) as non-linear activation functions (LeCun *et al*, 2015). The full architecture of the network is shown in Figure EV3B. The model was trained using Dice loss (Léger *et al*, 2018). The Adam optimization algorithm was used with a fixed learning rate of 10^-4^ (Kingma & Ba, 2014). The number of epochs (i.e., the number of cycles through the full training set) was chosen so that convergence was reached. The batch size (i.e., the number of images in every training iteration) was set at one.

For segmentation of the nucleoli, six U-net models were trained in parallel on the same set of training images, but considering different weight initializations. During the testing phase, segmentations were computed with all six models. A pixel was then classified as a part of the nucleolus if at least two predictions labeled it as such. This strategy increases the robustness of the approach. For segmentation of the nuclei, a single U-net model was used.

#### Learning strategy

The learning strategy is shown in Figure EV3C. Each DHM image was first normalized by dividing its pixels by their maximal intensity. Then, two separate deep learning models were trained. One segmented the nucleoli and the other segmented the nuclei. The nucleolus and nucleus segmentation masks automatically produced from the fluorescence images were used as labels during the training phase. For nucleolus segmentation, the binary masks obtained for estimating the nucleolar area were used. During the test phase, the processing pipeline analyzed only the DHM images, and the nucleolus and nucleus prediction binary masks were multiplied element-wise in order to remove potential false nucleoli detected outside the nuclei in the automatic nucleolus segmentation output (see ⊗ symbol on Figure EV3C).

We performed 3-fold cross-validation on the 75 images of the first dataset. This consisted in partitioning the 75 images into three subsets of 25 images. The model was trained on two subsets (i.e., on 50 images), with subset switching in order to test the model on all 75 images. This approach allowed computing the statistics on 75 images without reporting results on the training images. No model retraining was considered for segmentation of the 50 images of the second dataset. One of the three models used for predictions on the first dataset was arbitrarily chosen.

#### Postprocessing for statistical computation

To remove clustered cells and cells whose nucleus touched the border of the images, a postprocessing step was automatically performed, as such cells may bias the statistics. Clustered cells were detected from the binary masks used for nucleus segmentation. In those images, isolated cells have a convex elliptical nucleus, whereas clustered cells have overlapping nuclei, leading to non-elliptical and concave shapes. Hence, clustered cells are detected by identifying those concave shapes. This is done by computing the ratio of the area of the considered region to the area of its convex hull. If the ratio is smaller than 0.95, the aggregated nuclei are removed from the automatic nucleus segmentation output (and the corresponding nucleoli as well). This postprocessing was applied to the segmentation results obtained on both the fluorescence and DHM images.

#### Calculation of the nucleolar optical thickness

To compute the nucleolar optical thickness, we used the mean intensity of a fixed area. We used a 12-pixel area size throughout this work. Using a 50-pixel area size provided similar results (data not shown).

#### Production of nucleolar Cry2olig cell line

An mCherry-Cry2olig sequence was amplified from plasmid pHR-mCh-Cry2olig (Addgene #101222)(Shin *et al*, 2017), using 5’-GCATCACCACCATCACCATGCCTGCAGGCTCGAGATGGTGTCTAAAGGCGAGG -3’ and 5’-CGGGCCCTCTAGACTCGATCAGTCACGCATGTTGCAGG-3’ as primers. The PCR fragment was integrated into the pcDNA5/FRT/TO plasmid (Thermo Fisher) by InFusion Snap assembly (Takara). This plasmid was linearized with *Sbf* I and *Xho* I and a synthetic g-block (IDT) containing the NoLS of SF3B2 (Scott *et al*, 2010), and the RGG domain of LAF1 was inserted (Elbaum-Garfinkle *et al*, 2015). HEK293 Flp-In™ T-REx™ cells (Thermo Fisher) were co-transfected with the final plasmid and the pOG44 plasmid (Thermo Fisher), and stable integrated clones were selected with hygromycin.

#### FRAP analysis of the nucleolar Cry2olig cell line

Cells were seeded into Lab-Tek chambered coverglass slides (Thermo Fisher). Expression of the construct was induced by incubation with 1μg/ml doxycycline for 24 h. Induction of expression was confirmed by western blotting (data not shown). The cells were kept in the dark after induction of expression. Imaging was performed with a 63x/1.4 oil DIC objective (Plan-Apochromat, Zeiss) on a Zeiss Axio Observer.Z1 microscope driven by MetaMorph (MDS Analytical Technologies, Canada) and equipped with a Yokogawa spinning disk confocal head, an iLas multipoint FRAP module, an HQ2 CCD camera, a laser bench from Roper (405nm 100mW Vortran, 491nm 50mW Cobolt Calypso and 561nm 50mW Cobolt Jive), and a stage-top incubator system (Live Cell Instruments) providing stable 37°C and 5% CO_2_. FRAP images were acquired over 5 min in 500 ms intervals. A defined area was bleached by a 98% pulse of the 561 nm laser for 50 ms. Activation of Cry2olig oligomerization was achieved with a 500 ms 5% 488 nm laser pulse.

#### Quantification analysis of the nucleolar Cry2olig cell line

For optical density analysis, an 12-pixel area at the center of each nucleolus was marked and the mean intensity in the DHM phase images was calculated. The operator was blinded to the aggregation state of the cells. Three hundred eighty nucleoli were analyzed (173 liquid-state nucleoli and 207 gel-state nucleoli). Statistics: liquid state = 173 nucleoli analyzed (mean 156.5/STD 33.58/SEM 2.55); gel state = 207 nucleoli analyzed (mean 192.8/STD 28.88/SEM 2.01). Mann-Whitney test p<0.0001.

## ACKNOWLEDGMENTS

We thank Emilien Nicolas (ULB) for setting up the initial experiments. We thank Laurent Desmecht (Ovizio s.a) and Gilles Mordant (UCLouvain, statistics helpdesk) for useful advice on data analysis. Research in the Lab of D.L.J.L. was supported by the Belgian Fonds de la Recherche Scientifique (F.R.S./FNRS), Université libre de Bruxelles (ULB), BioWin [HOLOCANCER], EOS [CD-INFLADIS], Région Wallonne (SPW EER) Win4SpinOff [RIBOGENESIS], Internationale Brachet Stiftung, the COST actions EPITRAN (CA16120) and TRANSLACORE (CA21154), the European Joint Programme on Rare Diseases (EJP-RD) RiboEurope and DBAGeneCure. Jean Léger (Research Fellow), and Christiane Zorbas (Postdoctoral Researcher) were recipients of F.R.S./FNRS fellowships.

## EXPANDED VIEW FIGURE LEGENDS

**Figure EV1. Off-axis differential interferometer (QMOD) setup used in this work.**

Description of the beam path and microscope configuration used. The diagram illustrates the detailed beam path in our QMOD setup. A purposely built versatile “plug in” DHM adapter (QMOD, developed together with Ovizio s.a.) was connected between the lateral port of a Zeiss inverted microscope and a Retiga R3 camera (Qimaging) used for imaging the DHM phase and all fluorescence channels. The camera was driven from the MetaXpress (Molecular Devices) environment. Images were processed with OsOne (Ovizio). Digital holograms were recorded with an incoherent light source (**HAL** lamp) on an inverted microscope adjusted for proper Köhler illumination and coupled to a QMOD interferometer and CCD camera. A bandpass filter (**BP**) was used to increase the coherence of the light and obtain the partial coherence required for holography. The image forming light rays passing through the specimen (**S**) were captured with the microscope objective (**O**) and directed from microscope lens **L1** to the QMOD interferometer. In the QMOD, a diffraction grating **G** induced splitting of the incident light beam into a diffracted beam (**1**, reference) and a non-diffracted light beam (**2**, object beam). A second lens (**L2**) placed at focal distance from the grating G reshaped both the diffracted and non-diffracted beams into beams parallel to the optical axis. A wedge (**W**) inserted in the optical path of the object beam induced a slight shift of the images produced by the diffracted and non-diffracted light beams. **C** is a compensating optical module placed in the optical path of the non-diffracted light beam to compensate for the light shift introduced by W in the diffracted beam. The diffracted beam is then recombined with the object beam and focalized by means of objective lens **L3** on the recording plane of a CCD camera, where the hologram is recorded. LED, illumination; FF, fluorescent filter cube. Fluorescence imaging: excitation illumination is emitted by a light-emitting diode (**LED**) and is directed to the sample (**S**) through a fluorescence filter cube (**FF**). Fluorescence emission by the specimen is collected by the objective, passes through the filter cube and **L1**, enters the QMOD, and finally reaches the CCD camera.

**Figure EV2. DHM detection of the nucleolus in cell lines of various origins.**

The various cell lines indicated were observed by correlative fluorescence-DHM. Cells were stained with DAPI. An example of a perinucleolar chromatin ring, lining the nucleolus, is highlighted with an arrowhead in the HeLa-FBL-GFP panel for reference (see Fig 2B for details). Scale bar, 20 μm.

**Figure EV3. Image processing pipeline.**

**A** Segmentation pipeline of fluorescence images. The dynamic range of the GFP and DAPI images was normalized to [0,1]. GFP images were further standardized image-wise. Seeds were automatically positioned on the pixels with local maximum intensity in the GFP images. Starting from those seeds, region-growing algorithms were used with a constant threshold for nucleolus area estimation and an adaptive threshold for counting. *I_seed,j_* is the pixel intensity of seed j, *τ* the constant, and *α* a scalar between 0 and 1. Unrealistic holes remaining in segmented nucleolar masks were filled in a postprocessing step. After normalization, the DAPI images were filtered with a Gaussian kernel and an adaptive threshold was applied. The threshold was adapted for every DAPI image on the basis of the background and foreground (i.e., nucleus) pixel intensity distributions. Further postprocessing of the segmented nuclear masks included a morphological closing operation, removal of false detections, and a final morphological dilation operation. The symbol ⊗ denotes pixel-wise multiplication between binary masks to ensure that only nucleolar pixels contained within the nuclear area were considered to be *bona fide* nucleoli. The binary mask output of fluorescence image segmentation is illustrated (insets).

**B** Architecture of the 2D U-net convolutional neural network. Convolution operators are filters that extract image patterns useful for segmentation. The analysis path (left part) allows capturing context, whereas the synthesis path (right path) and skip connections allow retrieving high output resolution.

**C** Segmentation pipeline of DHM images. Two neural networks were trained, one with binary masks of nucleoli obtained by segmentation of the GFP images, the other with binary masks of nuclei obtained by segmentation of the DAPI channel. The binary mask output of DHM image segmentation is illustrated (insets).

**Figure EV4: Benchmarking of data analysis.**

**A** Conservation of the mean nucleolar area according to the number of nucleoli per cell. The predicted value is a theoretical upper limit, considering that the area occupied by a single spherical nucleolus is multiplied by N^1/3^ when the nucleolus is partitioned into N identical spheres.

**B** Histogram representing the number of nucleoli per nucleus automatically counted on the fluorescence images (left) and DHM images (right). Confusion matrices assessing automatic counting of nucleoli on the fluorescence images. The matrices compare the manual and automatic counts of nucleoli on 25 images. Element (i,j) in the confusion matrix is the number of nuclei with i nucleoli for which the algorithm counted j nucleoli. The elements on the main matrix diagonal were correctly counted. The lower the total value of elements outside the main diagonal, the better the counting.

**C** Sensitivity (true positive rate), specificity (true negative rate), and precision (proportion of properly classified cells) of the cell classification based on their number of nucleoli. Sensitivity provides the probability of counting *n* nucleoli amongst cells truly displaying *n* nucleoli. Specificity provides the probability of not counting *n* nucleoli amongst cells displaying *n* nucleoli. Precision provides the proportion of cells correctly classified. The closer the sensitivity, specificity, and precision are to 1, the better the approach.

**Figure EV5. Comparison of segmentation of the GFP signal by thresholding and of the DHM signal by deep learning.**

**A** Segmentation of the GFP signal by thresholding. Red, automatic segmentation of nucleoli on GFP images by ‘region growing’, for computing the nucleolar area; blue: automatic segmentation of nucleoli by ‘region growing’, for counting; yellow: manual annotation for validation of the nucleolus counting algorithm; green: automatic segmentation of the nuclei on GFP images by thresholding; gray, eliminated cells (image edges, clustered cells, etc.).

**B** Segmentation of the DHM signal by deep learning. Red, automatic segmentation of nucleoli on DHM images by deep learning (for counting, area computation, and nucleolar optical thickness computation); green, automatic segmentation of the nuclei on DHM images by deep learning; yellow, same as in panel A.

## SUPPLEMENTAL MOVIES

**Movie 1. DHM live-cell imaging through mitosis.**

Representative live imaging of HeLa-FBL-GFP cells. Cells were imaged with a 20x objective over a 12-h period in 1-min intervals in the DHM and fluorescence modes.

**Movie 2. DHM live-cell imaging upon roscovitine treatment.**

Live imaging of HeLa-FBL-GFP cells treated with 50 μM roscovitine in 0.1% DMSO. The red arrow indicates the first detectable changes in nucleolar morphology. Cells were imaged with a 20x objective over a 3-h period in 30-s intervals in the DHM and fluorescence modes.

